# LigHTS: Massively Parallel Biomimetic Photo-Functionalization for Imaging-Based Ultra-High-Throughput Screening

**DOI:** 10.1101/2025.10.23.683892

**Authors:** Alessandro Enrico, Sara Rigolli, Julius Zimmermann, Melissa Pezzotti, Eloisa Torchia, Moises Di Sante, Ferdinando Auricchio, Francesco S. Pasqualini

**Author notes:** A. Enrico and S. Rigolli contributed equally to this work.

## Abstract

Imaging-based ultra-high-throughput screening (UHTS) in pharma and biotech still runs on 384/1536-well plates whose stiff, flat substrates limit biological fidelity and screening efficiency. Highly biomimetic organs-on-chips and organoids improve relevance but lack reproducibility and plate-scale throughput. Biomimetic hydrogel scaffolds can be produced at scale through photopolymerization, which yet uses focused optics to define micrometer-resolved geometries, constraining scalability. To address the technical challenge of truly scalable biomimetic substrates featuring anisotropies, this study presents LigHTS, an all-optical, in-well method that replaces focused with collimated illumination to photofabricate structured hydrogels in standard 384/1536-well plates. Adding food dye tartrazine to gelatin-methacrylate (GelMA) solutions increases hydrogel thickness sensitivity to UV dose by ∼10x, allowing uniform control of film thickness without lenses. Entire plates are functionalized in parallel with soft hydrogels (∼1-10 kPa) whose thickness is tunable from 10 to 100 µm. Simultaneously, simply interposing film photomasks encoding anisotropies enables orthogonal control of thickness and topography at UHTS throughput. Biological effect is demonstrated with mechanosensitive HT1080 cells, which display stiffness- and topography-dependent spreading and contact-guided migration on LigHTS-produced grooved substrates. Geometric uniformity across the plate (coefficient of variation <20%) meets HTS reproducibility standards, providing a readily available solution with enhanced biomimicry for imaging-based UTHS pipelines.

## 1. Introduction

Increasing the efficiency of drug discovery and development is crucial for the pharmaceutical industry and society.^1–3^ Despite many efforts to streamline the pipeline, this process remains extremely costly and risky, taking up to 15 years and more than $1 billion on average per approved new drug.^4,5^ One contributor to this high failure rate is the poor predictivity of early-stage drug screening models that lack biomimicry.^6,7^ To remain economically viable, even modern imaging-based ultra-high-throughput screening (UHTS) still relies on conventional ultra-high-throughput formats, mainly 384-/1536-multiwell plates. A typical drug screening campaign tests ∼10,000 compounds and demands costs of only a few $ cents per datapoint, therefore inexpensive 384-or 1536-well plate assays are the golden standard despite their lack of biomimicry.^8^ In fact, in these formats cells are still cultured on stiff, flat substrates, which are convenient for imaging automation and analysis, but far from the structured and mechanically compliant microenvironment of real tissue.^9,10^ We argue that this is at least in part why many drug candidates that seem promising in conventional HTS ultimately fail in vivo (false positives), leading to expensive late-stage failures.^11^ At the same time, oversimplified in vitro models can yield false negatives, causing valuable drug candidates to be erroneously discarded and only repurposed decades later.^12^

Biomimetic models such as organoids and organ-on-chips indicate that recreating native tissue conditions can improve the predictive accuracy of preclinical assays.^13–15^ Organoids are self-assembled 3D cell aggregates that recapitulate key aspects of tissue architecture and can be produced in relatively high throughput. However, their inherent variability clashes with the strict reproducibility requirements of UHTS in terms of coefficient of variation (±20% CV).^16,17^ Their 3D nature further complicates automated imaging and analysis, ultimately limiting their use in early-stage screening.^18,19^ Organ-on-chip systems, by contrast, reliably emulate tissue microenvironments in microfluidic devices (with perfusable channels and organ-specific compartments) and often integrate sensors for real-time and high-content readouts.^20,21^ Yet these sophisticated platforms come at a high cost (up to $1000 per device), making them incompatible for large-scale studies.^22–24^

To bridge the gap between biomimicry and UHTS requirements, a cost-effective way to incorporate a biomimetic scaffold into UHTS formats is needed.^25^ Ideally, this solution would provide a compliant hydrogel scaffold with physiological stiffness (∼1-10 kPa for soft tissue) in each well.^26^ The hydrogels should be in the optimal thickness range (10-100 µm) to isolate the cells from the rigid substrate, minimize spherical aberration and scattering that are detrimental to imaging, and remain within the working distance of high-resolution objectives (around 370 µm, effective <150 µm of dynamic considering the ∼200-µm-thick glass substrate).^27^

Ideally, the hydrogel surface would also carry anisotropic features that can provide cell- and tissue-organizing cues, such as microgrooves, that are important for recapitulating cell phenotypes in structured microenvironment, such as in cell migration.^28–30^ Hydrogel surface structuring can be made via micromolding on flat substrates in low throughput, but to our knowledge no published method or product offers this functionality in standard 384-or 1536-well plates. We previously developed a process to functionalize plates up to the 384-well format with thin, meniscus-free, high-resolution imaging-compatible fish gelatin hydrogels using automated liquid handling, with the caveats of incomplete well coverage and lack of hydrogel surface structures.^31^ On the other hand, there are commercially available formats up to 384-well plates featuring hydrogel scaffolds with stiffness and complete well coverage, which also lack surface structures and are in addition too thick (>100 µm) for optimal imaging, especially at high resolution and in low signal-to-noise conditions.^32,33^ Recent published works have reported partially biomimetic functionalization on 96-well plate formats, using either molded/embossed polymer topographies integrated into the plate base,^34–40^ mostly relying on the fabrication of personalized plates starting from glass slides and bottomless plates.^41–43^ These techniques show strong prototyping potential in academic contexts, but scaling hydrogel manufacturing to 384/1536-well plates with +/-20% CV for the critical geometrical parameters with ±20% and a <1$ cost per well is an open challenge.

In principle, optical patterning would be the most promising route to meet UHTS constraints because it is intrinsically non-contact and potentially scalable. However, current approaches are limited to surface sculpting and still rely on molding-based thickness definition.^44–46^ In situ spacer-free photopatterning requires a precise control of the axial cure depth which is complicated by oxygen inhibition, light scattering, and well-to-well volume non-uniformity, problems that are exacerbated in small-well geometries.^47^ Similarly, to obtain highly reproducible surface structuring, it is necessary to develop ways to translate spatially structured UV exposure into surface features in the depth of the hydrogel precursor solution. Even with a state-of-the-art digital micromirror device (DMD) or other projection optics, the limited exposure area/the use of focused lenses causes stitching errors which limit the uniformity of hydrogel thickness and mechanical modulus as well as groove fidelity to CV value above the UHTS tolerance across plates, and the serial nature of focused illumination prevents process scalability.^48^

To solve these problems concurrently, we took inspiration from conventional 3D printing, and we reasoned that the inclusion of a strong UV absorber competitor could dominate the light absorption dynamics, effectively decoupling it from the concentration and dynamic bleaching of the photoinitiator.^47,49,50^ Using light-matter interaction to control micrometric geometry removes the need for focusing optics and the associated limited throughput. We hypothesized that widefield or collimated UV light could be used for plate wide exposure with processing speed and fabrication costs far exceeding conventional 3D printing serial approaches, making them compatible with UHTS.

Here, we report on LigHTS, an all-optical scalable method to form microgrooved GelMA hydrogels directly inside 384- and 1536-well plates (**Figure 1a**). Introducing a biocompatible strong UV absorber, tartrazine, in a hydrogel precursor solution allows to define meniscus-free hydrogels in the 10-100-µm thickness range without the need for focusing optics (**Figure 1b).** Plate-wide collimated exposure enables stitching-free fabrication with ±20% CV across key geometric features and low per-well cost (<1 $). Uniform illumination can be structured by interposing photomasks during the hydrogel UV curing step to obtain surface structures (parallel microgrooves in this study) with single-digit-micron geometrical fidelity. The resulting hydrogels are thickness- and properties-compatible with low- and high-resolution fluorescence imaging as well as holographic phase imaging, making LigHTS extremely relevant for imaging-based phenotypic screening. As a proof of concept, we seeded HT1080 human fibrosarcoma cells with genetically encoded fluorescent proteins on LigHTS hydrogels to study the modulation of the static (cell area) and dynamic (topography-guided migration) phenotype due to soft, surface-structured substrate (**Figure 1c**).^51^ LigHTS thus combines physiological stiffness with anisotropic architecture in UHTS formats to improve the predictive value of imaging-based screening.

**Figure 1.**
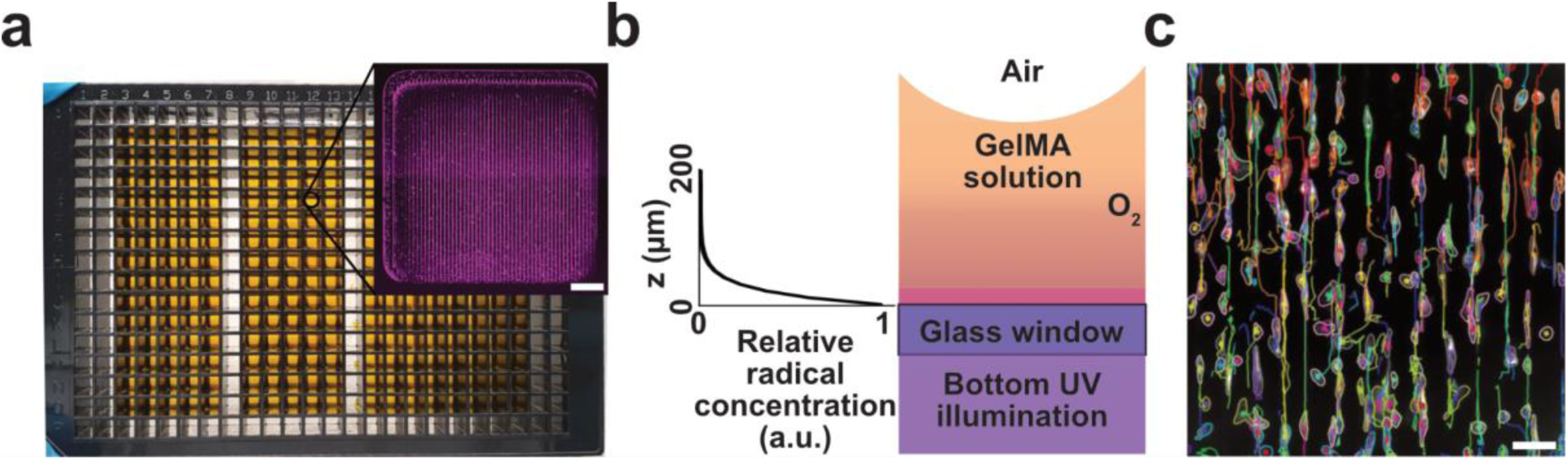
LigHTS: one-step, light-patterned, solution bringing soft grooved gels to high-throughput screening. a) UHTS-compatible fabrication showcased with gel patterning in a 384-wellplate format. The same single UV exposure (16 minutes) can simultaneously pattern thickness-controlled (<100 µm) stiffness-compliant (E ≈ 1-10 kPa) grooved gels on the well bottom of UHTS vessels. The method is compatible with the infrastructure and price targets of pharmaceutical HTS pipelines, and requires no clean-room lithography, integration, or additional gluing steps post patterning. Scale bar in the inset, 500 µm. b) Process principle. A soluble UV-absorbing chromophore is added to a GelMA prepolymer. Following Beer-Lambert attenuation, the chromophore creates a steep, exponential decay in photon flux, and thus radical generated by photoinitiator cleavage. As a result, hydrogel thickness can be controlled by tuning the UV dose, UV absorber concentration, and GelMA content. c) Imaging-based phenotypic screening of cells in a 2.5D biomimetic environment. In the presence of grooved gels, HT1080 cell lines display contact-guidance migration with a preference for the groove main axis. Scale bar 100 µm.

## 2. Results

Our method relies on doping of photocurable solutions with a strong optical absorber to obtain sub-100-µm thick hydrogels by non-focused UV illumination. In essence, hydrogel precursor solutions were cast in glass-bottom multiwell plates and later bottom illuminated to induce chain photopolymerization (Figure 1b, see Experimental section). Multichannel pipettes can be manually or robotically operated to deliver sufficient volume in each well to ensure complete glass substrate wetting (typically >20 µL for 384-well plates and >4 µL for 1536-well plates). To systematically interrogate the parameter space, we performed serial exposures of 384-well plates using a DMD and varying UV dose (3 to 21 mJ mm^−2^), as well exploring the effect of different GelMA (50 to 200 mg mL⁻¹) and tartrazine content (up to 9 mg mL^−1^, see Experimental section).

To study the behavior of conventional hydrogel precursor solution in this configuration as baseline, we first utilized a commercially available solution of GelMA (75 mg mL^−1^ in 1:10 v/v glycerol:PBS) and lithium phenyl-2,4,6-trimethylbenzoylphosphinate photoinitiator (LAP, 2.5 mg mL^−1^, **Figure 2a**, see Experimental section). For all tested doses, we observed complete volumetric curing of the hydrogel solution present in the wells. We then coated the resulting hydrogels with fluorescent beads and used confocal z-stacking to precisely evaluate gel thickness and hydrogel surface profile (see Experimental section). The volume reconstruction shows that the UV curing produced millimetric-thick hydrogels freezing the high-curvature air-liquid interface (in-well height residuals exceed 200 µm, **Figure 2b-c**), making these hydrogels incompatible with high-resolution imaging and sub-optimal for cell culture uniformity.

**Figure 2.**
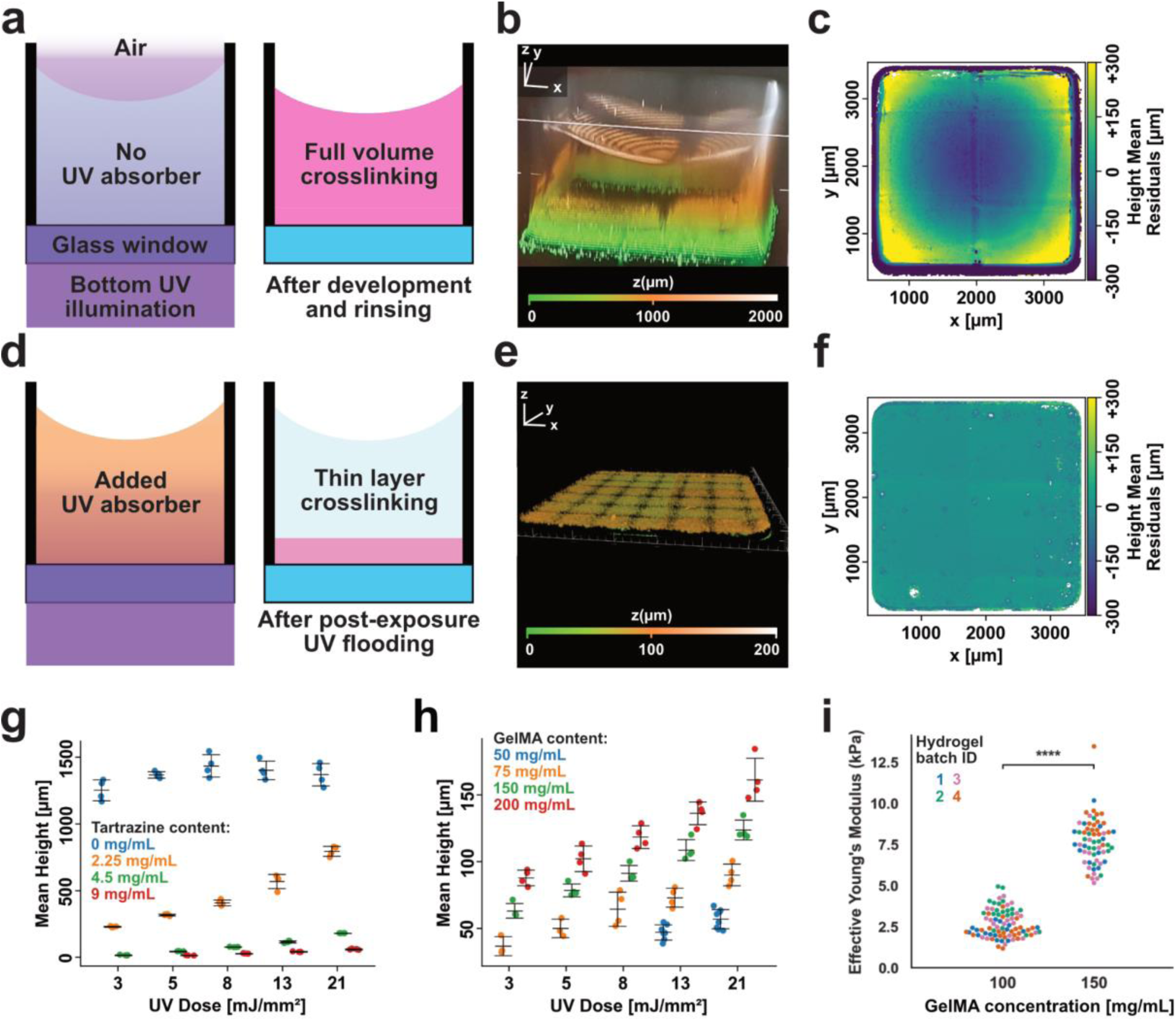
Introducing strong optical absorber in hydrogel precursor solution enables sub-100-μm thickness-tunable flat gels in non-focused UV exposure configuration. a) Polymerization dynamics for conventional hydrogel precursor solution in wellplate formats. When illuminated with a sufficient UV dose, the entire prepolymer solution crosslinks. **b)** 3D isometric rendering and **c)** 2D residual height map of resulting hydrogels with prominent meniscus (>200 μm difference in gel thickness between center and sidewalls). The resulting thickness is also strictly above 100 μm, mainly determined by the available volume of solution, with resulting scattering. The hydrogel surface was coated with fluorescent nanoparticles and analyzed with confocal fluorescent microscopy to reconstruct the hydrogel profile. **d)** LigHTS strategy consisting in doping photosensitive hydrogel precursor solution with a strong UV absorber, such as tartrazine, to decouple optical absorption from the ECM and photoinitiator components and enable single-digit-micrometer control of the hydrogel thickness. **e)** 3D isometric rendering and **f)** 2D residual height map of meniscus-free, flat (10-fold decrease in thickness inhomogeneity), and thin hydrogels obtained by ligHTS approach. **g)** Parametric study of thickness dependence on UV dose and tartrazine concentration for fixed GelMA (75 mg mL⁻¹) and LAP (2.5 mg mL⁻¹) concentration. Hydrogel thickness decreases monotonically with tartrazine concentration and increases with UV dose, as expected (N=4 gel per condition). **h)** Parametric study of thickness dependence on UV dose and GelMA concentration for fixed tartrazine (4.5 mg mL⁻¹) and LAP (1 mg mL⁻¹) concentration. At each given dose, increasing GelMA content increases the resulting hydrogel thickness (N=4 gel per condition). **i)** Programmable compliance measured by nanoindentation experiments. Each datapoint refers to the extracted Young modulus from an indentation curve using the Hertzian contact model. At least ten measurements were performed on each replicate per experiment (N=4). Increasing GelMA concentration tunes the modulus from around 2.5 kPa (100 mg mL⁻¹) to around 8 kPa (150 mg mL⁻¹). Significance bars and stars between groups represent statistical significance for pairwise test (p<0.0001).

To reduce the thickness of the resulting gels and increase the thickness sensitivity to UV dose, we included tartrazine, a food dye absorber, to act as non-reacting, non-bleaching competitor to LAP for photon absorption (**Figure 2d**). We first characterize the molar extinction coefficient at 365 nm from LAP (around 220 M^−1^ cm^−1^) and tartrazine (8000 M^−1^ cm^−1^), finding that tartrazine has a significantly higher (>10x) absorption than LAP at 365 nm and stable transmission, suggesting no significant bleaching. Mixing tartrazine in the commercial hydrogel precursor solution radically changes the curing behavior: the resulting optical absorption is significantly stronger than the one in the base hydrogel precursor solution, and light does not penetrate through the bulk of the liquid. As a result, the production of radicals matches the exponentially decreasing profile in the z direction. In these chemical systems, dissolved oxygen is quenching photogenerated radicals faster than their interaction with GelMA, and sol-gel conversion effectively occurs after oxygen is locally depleted (oxygen burn-in). In tartrazine-containing solution, the density and radical rate production decay exponentially with propagation depth through the hydrogel precursor solution (Figure 1b), resulting in through-thickness crosslinking gradient due to optical attenuation and oxygen inhibition. Moreover, the presence of oxygen is ultimately hard capping hydrogel thickness regardless of the UV dose, given solution composition and light intensity, with lower methacrylate conversion at depth. After removing tartrazine in the development steps (see Experimental section), a post-exposure UV flooding in a LAP-containing PBS produced homogeneously crosslinked gels across the thickness (Figure 2d).

To quantitatively characterize hydrogel thickness and flatness, we performed confocal z-stacking by manually verifying the position of the glass-hydrogel interface and developed a custom-made analysis pipeline in Python (see Experimental section). The hydrogels produced with the LigHTS method are significantly thinner and flatter than the ones from the base hydrogel precursor solution (**Figure 2e** and **f**). We found that the inclusion of the UV absorber significantly (p<0.001) decreases the thickness of the resulting gels at every tested UV dose, and allows precise control of gel thickness. The resulting hydrogel thickness decreases 4-, 20- to 50-folds for tartrazine contents of 2.25 mg mL⁻¹, 4.5 mg mL⁻¹, and 9 mg mL⁻¹, respectively **(Figure 2g)**. At 9 mg mL⁻¹ tartrazine and 3 mJ mm⁻² dose, it was not possible to characterize the hydrogel thickness, indicating either no gel or very shallow (<5 µm) ones, while at 4.5 mg mL⁻¹ tartrazine and 21 mJ mm⁻² dose, the hydrogels exceeded the z-stack range (>180 µm), outside the range of interest for high-resolution microscopy. In presence of tartrazine, the gels display high uniformity/flatness (<10% thickness variation over the entire well, and standard deviation in the single-digit-micron range). Moreover, gel thickness displays a 10-fold increased sensitivity to the UV dose within the tested range, enabling fine thickness tuning by dose adjustment. The absolute thickness sensitivity to UV dose is monotonically decreasing with increasing tartrazine, resulting in higher dose and thus exposure time to achieve the same gel thickness. While both 4.5 mg mL⁻¹ and 9 mg mL⁻¹ tartrazine-containing formulations can produce hydrogels in the 10-100 µm thickness range, using 4.5 mg mL⁻¹ tartrazine requires four times lower dose/exposure to achieve the same gel thickness as compared to using 9 mg mL⁻¹. The 4.5 mg mL⁻¹ tartrazine concentration therefore strikes the best balance between relative sensitivity for thickness tuning and absolute sensitivity for time-efficient patterning.

To modulate the stiffness of the hydrogels, we fixed the tartrazine concentration to 4.5 mg mL⁻¹ and switched from ready-made low-GelMA solutions to custom-made ones with a GelMA concentration going from 50 to 200 mg mL⁻¹ (**Figure 2h**, see Experimental section). LAP concentration was reduced to 1 mg mL⁻¹ to increase the thickness tuning range while using the high UV intensity of the DMD illumination system. Increasing GelMA content results in a significant monotonical increase in gel thickness. All hydrogel formulation resulted in hydrogels upon 3 and 21 mJ mm⁻² UV illumination, except for the 50 mg mL⁻¹ which did not form any hydrogels for the 3 and 5 mJ mm⁻².

To verify that the gels obtained with this method followed the same trend, we used microindentation experiments to assess the stiffness of the gel surface after UV flooding (**Figure 2i**, see Experimental section). We selected 5 µm indentations with a cell-sized (25-µm-diameter) probes to assess the conditions cells would effectively interrogate compared to the bulk-property results from rheology or discrete stiffness level of the hydrogels components characterized via atomic force microscopy. We obtained stiffness values in the range of interest for soft tissue (1-10 kPa) for different GelMA content (100 and 150 mg mL⁻¹), in line with the literature. These results confirm that surface stiffness still scales with local conversion and polymer volume fraction, despite the presence of tartrazine and inhomogeneous crosslinking before the UV flooding.

To study the compatibility of the resulting hydrogels after tartrazine removal and UV flooding with high-resolution microscopy, we perform high resolution live-cell imaging acquisition in confocal mode using a 100× silicone oil objective (see Experimental section), and HT1080 human fibrosarcoma cells expressing fluorescent markers for actin and tubulin actin we recently engineered (see Experimental section).^51,52^ Compared to the standard of cells directly seeding on glass substrate (**Figure 3a**), we could obtain high-quality images on hydrogels with thicknesses ranging from 20 to 80 µm (**Figure 3b** and **b**), indicating high optical clarity of the hydrogels, negligible fluorescence background, and minimal spherical aberration resolution loss in the hydrogel thickness range of interest. To characterize the impact of cell substrate stiffness on the static cell phenotype, we segmented the cells in 20x acquisitions combining actin and tubulin signals and compared the resulting cell area (see Experimental section). We found that the area was decreasing with decreasing substrate stiffness (**Figure 1e**), showing significant (p<0.001) modulation of the cell phenotype even between softer and stiffer gels within the 1-10 kPa range.

**Figure 3.**
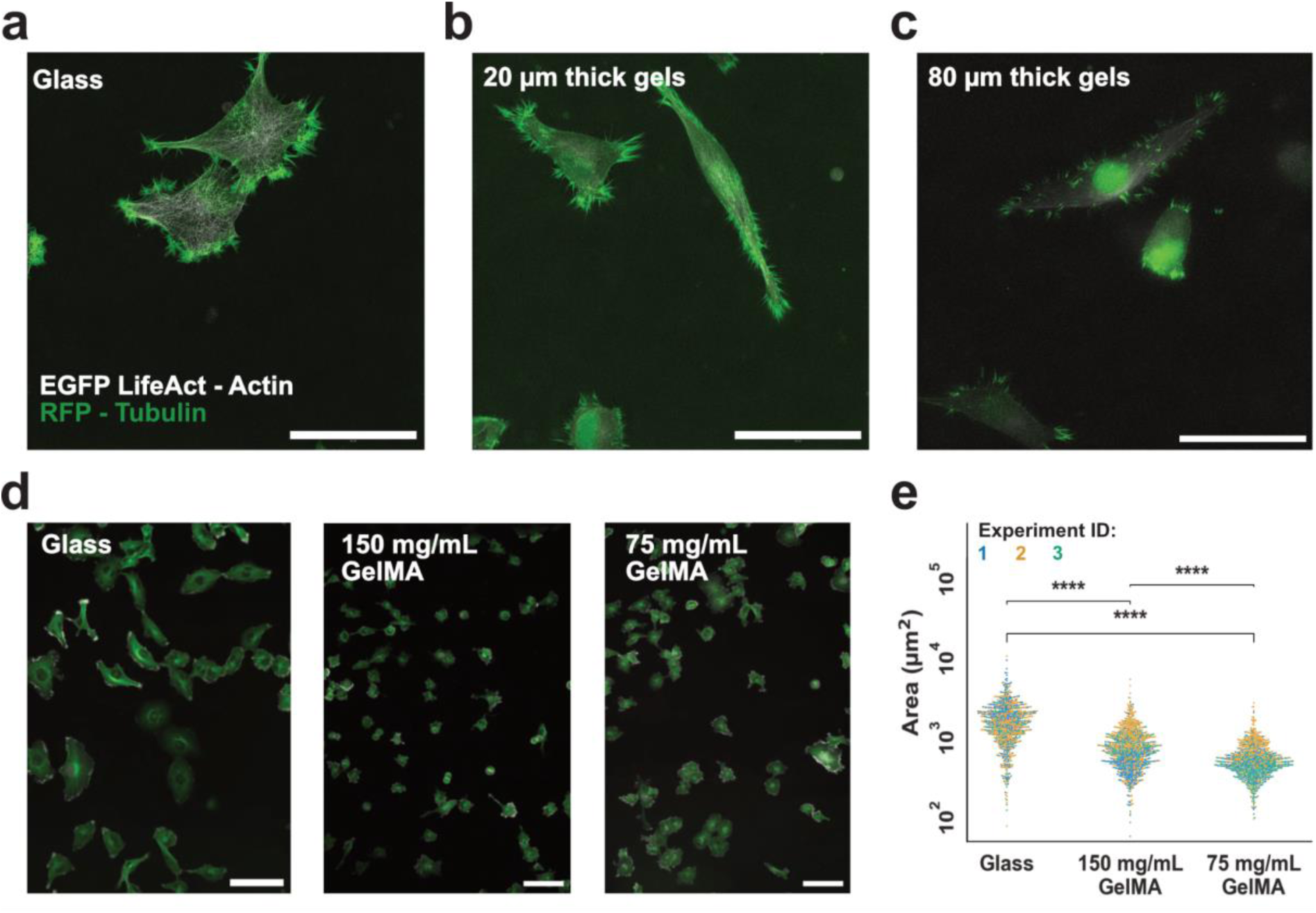
LigHTS hydrogels couple tissue-like mechanics with microscope-friendly optics, enabling high-content phenotypic screens. a-c) Fluorescence confocal imaging of HT1080 fibrosarcoma cells stably expressing Actin-EGFP/Tubulin-RFP on glass (panel a) and on thin soft (panels b and c) hydrogels at high (100x, silicon oil immersion objective) magnification. Because LigHTS films are ≤100 µm thick and sit on the coverslip, they remain within the working distance of high-numerical-aperture objectives. Sub-cellular features are crisply resolved on a) glass and on LigHTS gels of b) 20 µm and c) 80 µm thickness, validating compatibility with high-resolution, single-cell read-outs. Nuclear signal comes from the bleedthrough of the engineered FUCCIPLEX cell cycle sensor for cells in G0/G1 phase. Scale bars, 20 µm in a), b) and c). d) Fluorescence confocal imaging of HT1080 fibrosarcoma cells at low (20x air immersion objective) resolution. 75 mg mL⁻¹ and 150 mg mL⁻¹ GelMA correspond to elastic moduli of 2.5 kPa and 8 kPa moduli. Scale bars, f) Characterization of HT1080 cell area on different substrates. Tubulin channel is used to extract cell area masks via Cellpose-SEM algorithm to demonstrate cell area modulation on substrates with different stiffness. Each point corresponds to a different cell (n > 150 per condition in each experiment), each color to a different experiment (N=3). Significance bars and stars between groups represent statistical significance for pairwise test (p<0.0001).

To simultaneously obtain hydrogels with controlled thickness in the 10-100-µm range and micrometric-surface structures without the need of pre-formed micromolded hydrogels, we used structured illumination (**Figure 4a**). In this study, we aimed for the generation of aligned microgrooves since such surface structures are relevant for cell polarization and tissue alignment, and targeted groove peak-to-peak pitch of roughly 21, 29, and 40 µm (**Figure 4b**). The specific values were selected to match the cell characteristic size and to provide pixel-perfect maps for structured illumination given pixel quantization in the DMD. To maximize the height difference between groove peaks and bottoms, we structure the illumination to feature sharp discontinuities meant to mitigate the feature blurring due to light scattering and diffusion of radicals (Figure 4a).

**Figure 4.**
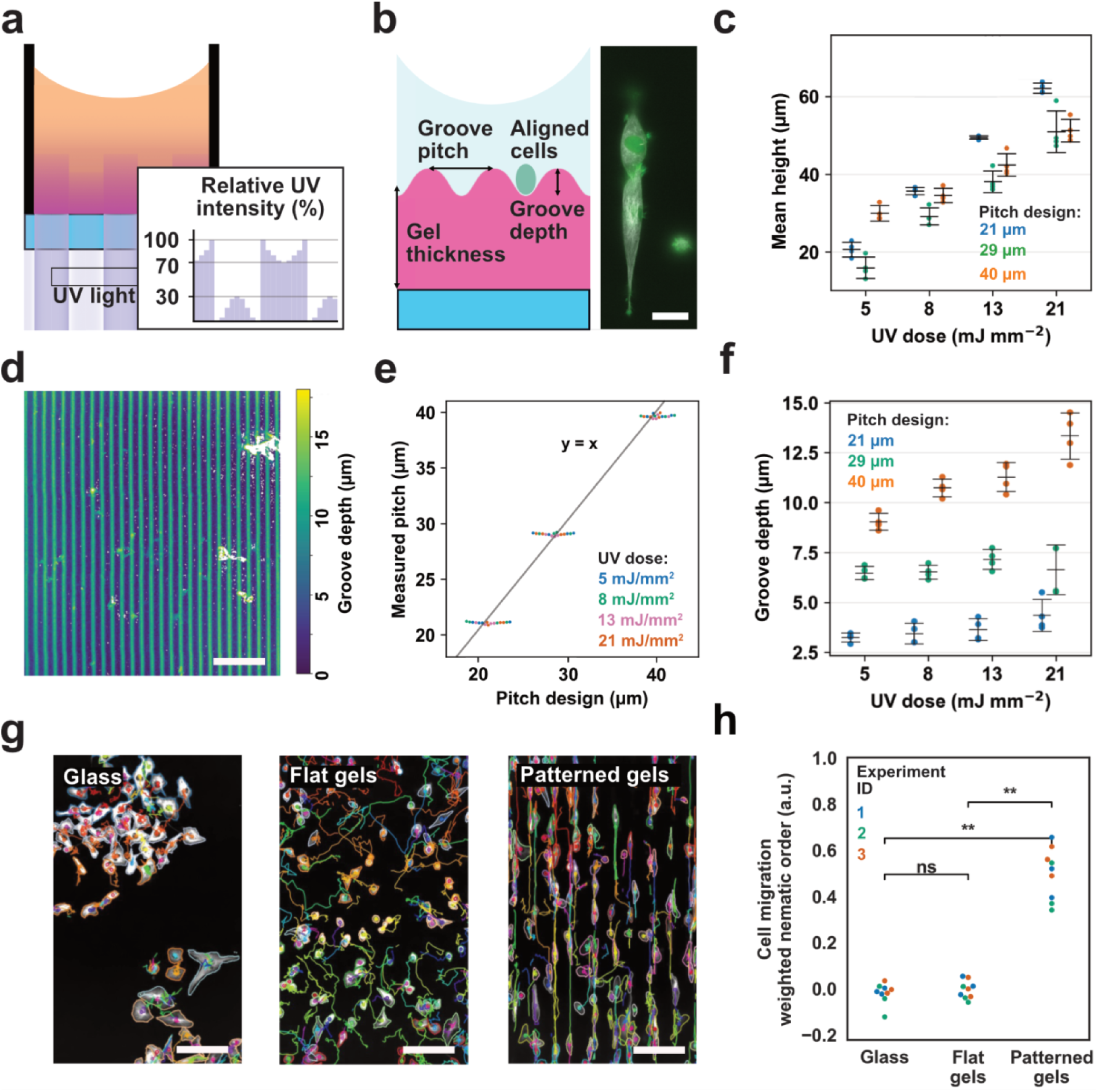
LigHTS fabrication method allows for surface photostructuring of hydrogels with single exposure, enabling contact guidance of cells. a) Gradient light intensity profile translates into surface photostructuring via a single non-focused UV exposure. b) Schematic illustration of the critical parameters in the grooved hydrogels, resulting in HT1080 cell polarization and maintaining high resolution compatibility. Scale bar, 25 µm. c) Mean hydrogel thickness of surface photostructured hydrogels with different UV doses and pitches. Gel thickness is mostly determined by the UV dose, with the pitch design slightly influencing the hydrogel thickness (N=4 per UV dose × pitch design). d) Heatmap reconstruction of photostructured hydrogel surface with groove depth quantification. Scale bar 200 µm. e) Measured pitch is highly conformal to the design across different spacings and different UV doses (N=4 per UV dose × pitch design). f) Quantification of groove depth as a function of UV dose for different pitches designs. Groove pitch is the dominant factor over UV dose in determining the groove depth (N=4 per UV dose × pitch design). g) HT1080 fibrosarcoma cell line migration experiment over a 12h span. Compared to glass and isotropic gels, surface-grooved hydrogels direct cell migration primarily along the main axis of the grooves. Scale bars 100 μm. h) Weighted Nematic Order parameter quantification of cell migration alignment (N=3 experiments, n=3 replicates per condition). Photostructured grooved contact-guide cell with statistical significance (p < 0.01) to migrate aligned with the groove axis with respect to isotropic substrate.

To explore the parametric space in terms of pitch design, we used solutions containing 150 mg mL⁻¹ GelMA, 4.5 mg mL⁻¹ tartrazine and 1 mg mL⁻¹ LAP. The resulting hydrogel precursor solution were illuminated, selecting nominal UV doses from 5 to 21 mJ mm⁻² then modulated according to the illumination patterned, resulting in a 2-fold decrease in large-area effective dose (**Figure 4c**, see Experimental section). As a consequence, the thickness values of the grooved hydrogels obtained with this approach were roughly half compared to the ones obtained with uniform illumination at any given doses (Figure 2h) and in the ideal range for mechanically-compliant HR-imaging-grade cell-scaffolds. To quantitatively characterize the resulting surface geometry and the reliability of the hydrogel photopatterning, we developed a custom-made analysis pipeline in Python to reconstruct and evaluate the hydrogel surface (**Figure 4d**, see Experimental section). In this study, we limited the analysis to Fields of View (FOVs) within the DMD printing fields, since stitching areas feature single-digit micron steps in height that are unrelated to the dynamics of the chemophysical dynamics of the system. The structured illumination produced grooves on the hydrogel surfaces with excellent matching between pitch design and measurement pitch with deviations in below the minimum lateral resolution of 20x acquisition (**Figure 4e**). In terms of groove depth, we observed the formation of peak-to-bottom height difference ranging from 3 to 15 µm (**Figure 4f**). Interestingly, the resulting groove depth was mainly modulated by the pitch design rather than the UV dose, which only significantly affected groove depths for the 40 µm design.

Finally, to characterize the impact of cell substrate stiffness and surface structure on the dynamic cell phenotype, we performed 12-hour migration assays for HT1080 cells seeded on patterned (40 µm pitch) and flat hydrogels as well as on glass, and we developed a dedicated analysis pipeline in Python for segmentation and tracking (**Figure 4g**). Specifically, we quantified the degree of migration aligned to the groove main axis irrespective of the verse, and therefore define as metric the displacement-weighted nematic order (WNO, assuming value of 1 for purely vertical direction, -1 for purely horizontal, and 0 for random migration directions, see Experimental section). Cells on grooved hydrogels had their migration significantly (p<0.01) polarized along the groove main axis, while cells on flat gels featured on average random migration direction (WNO around 0, **Figure 4h**).

Having characterized the process capability and explored the parametric space for both flat and grooved hydrogels, we transitioned from prototype-friendly yet serial DMD structured illumination to a uniform 10 cm × 10 cm illumination with a collimated UV source to achieve massively parallel patterning. Specifically, we used highly collimated light from a conventional mask aligner and illuminated the plate upside down **(Figure 5a).** This configuration enables the simultaneous exposure of an entire multiwell plate. The combination between room temperature patterning and the 10% v/v glycerol content in the hydrogel precursor solution prevents leaking of the solution in this plate-inverted configuration. To obtain grooved hydrogels, we considered interposing a photomask with parallel opaque lines with resulting groove pitch of 40, 60, and 80 μm to obtain a structure illumination. We studied how diffraction would transform the illumination profile within the hydrogels (**Figure 5b**), since the mask cannot be placed in contact with the hydrogel and the minimum distance from the glass-hydrogel interface is equal or greater than the glass substrate thickness (typically around 180 μm). Due to diffraction, the digital illumination at the mask level is transformed to a sinusoidal pattern at the hydrogel level, conveniently allowing simultaneous thickness and surface structure definition. Interestingly, even lower collimated light can produce suitable light patterns, enabling photopatterning with this scale and precision using low-cost exposure setup.

**Figure 5.**
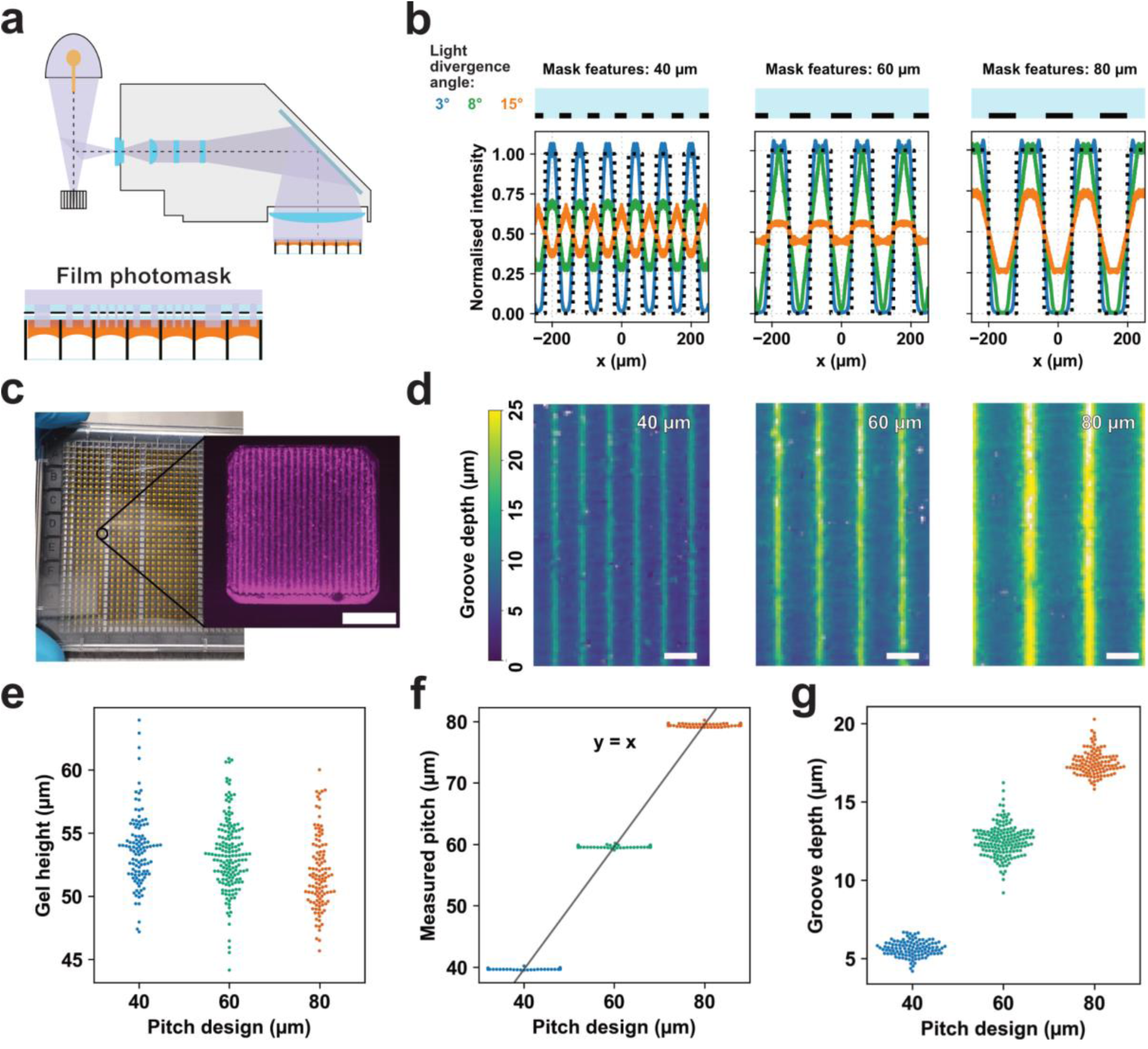
Ultra-high-throughput parallel fabrication of surface photostructured hydrogels. a) Schematic of the collimated UV-light system used for parallel whole plate fabrication through a film photomask. b) Simulated light intensity profiles across masks with different pitches, illustrating the effect of light divergence angles on hydrogel structuring of grooves. c) Proof of concept of parallel hydrogel photostructuration in standard 1536-well plate format, with inset showcasing patterned grooves. Scale bar 500 μm. d) Heightmap reconstructions of photostructured hydrogel surfaces through photomask with different pitches showcasing groove depth quantification. Scale bar 200 µm. e) Mean hydrogel thickness after 16 minutes UV exposure. The graph shows good uniformity between the same design, with a slightly yet statistically significant decreasing trend at increasing pitch spacing (N>100 wells per pitch). f) Measured pitch is highly conformal to the design across different spacings (N>100 wells per pitch). g) Groove depth is consistent between different replicates exposed to the same design, while pitch spacing showcases a statistically significant increase in groove depth (p<0.0001, N>100 wells per pitch).

To test the scalability of this process, we performed parallel patterning of all the hydrogels contained in the wells for 384- and 1536-well plates in a single 16-minutes UV exposure at roughly 3.5 mW cm^−2^ **(Figure 5c)**. Using the same characterization pipeline developed for serial photopatterning of grooved gels, we successfully demonstrated that we could bring continuous and well-defined microgroove structures in UHTS formats **(Figure 5d)**. All hydrogels formed with a bulk thickness between 50 and 60 μm, excellent groove pitch fidelity **(Figure 5e)**, and groove depth monotonically increasing with groove pitch designs, ranging 7 μm at 40 μm pitch to 17 μm at 80 μm pitch (p < 0.0001, **Figure 5g**).

To test the compatibility of this process with industrial requirements, we quantified the in-plate coefficient of variation (CV) for geometrical features, demonstrating that resulting hydrogel thickness and groove pitch are within ±20% CV range (N>300 for 1536-well plate, **Figure 6a**), with the pitch fidelity showing even higher reproducibility (±3% CV, **Figure 6b**). The resulting groove depth spread is within the 20% CV range, partly due to the limitation of the quantification method (measurement error for the z-evaluation in 20x confocal acquisition, **Figure 6c**). To complete the validation of whole-plate patterning against DMD-produced structures, we performed the same migration assay, obtained a similarly polarized cell migration along the groove main axis (**Figure 6d**). Cell migration is polarized by all the different patterns with high internal reproducibility (±20% CV). Interestingly, statistically significant differences emerged between different pitches, with 60 μm pitch showing the best results in directing cell migration (p < 0.0001 in pairwise tests against the other pitches).

**Figure 6.**
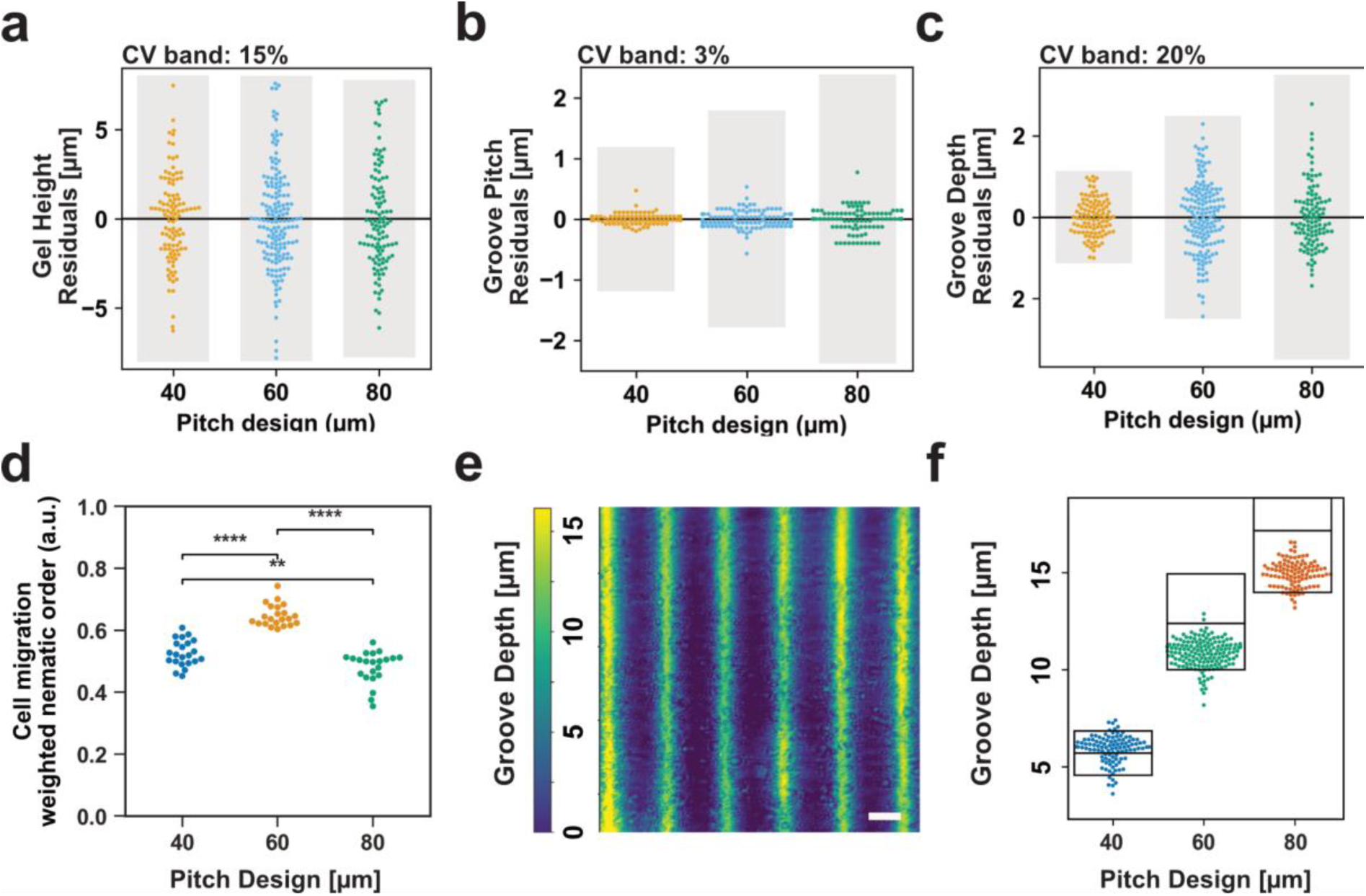
Reproducibility of LigHTS parallel photostructuring and validation by Holographic Phase Imaging. a-c) LigHTS-produced hydrogels show reproducibility up to industry-grade standard (N>100 wells per pitch), with a coefficient of variation under 20% for gel thickness (panel a), under 3% for pitch design compliance (panel b), and under 20% for grooves depth (panel c). d) WNO quantification of HT1080 cell line migrating along parallel photostructured gels with different pitches (N=20 wells per pitch). Statistical analysis showcases significative difference in migration alignment between 60 µm pitches and the other groups (p<0.0001) and a smaller but still significant difference between 40 and 80 µm (p<0.01) e) Heatmap showcasing grooved depth quantification via Holographic Phase Imaging. Scale bar 50 µm f) Quantification of grooves depth via Holographic Phase Imaging (N>100 wells per pitch). Boxes represent average groove depth and +/-20% CV band characterized with confocal fluorescent z-stacking. A good accordance is shown for wells with the same pitch, while increasing the pitches also increases groove depth as previously showcased. With respect to confocal analysis, Holographic Phase Imaging tends to underestimate groove depth, as shown by black boxes, representing depth average and +/-20% CV bands of the matching confocal characterization.

To speed up the quality control and quantitative evaluation of non-fluorescently tagged hydrogels and cells, we performed holographic phase imaging on the grooved hydrogels. Holographic phase imaging requires significantly lower exposure time (typically 1-10 ms) than confocal microscopy (typically 0.1-1 s), providing a potentially scalable characterization method that could be effectively implemented in pharmaceutical pipelines. We successfully demonstrated that this technique can be used to qualitatively reconstruct the surface profile (**Figure 6e**), quantitatively perform quality control of the surface microstructure (**Figure 6f**), and tracking cell migration. Compared with the groove depth characterization on the confocal microscopy dataset (Figure 5g), there is a substantial quantitative agreement between the values of the two different characterization methods. These results confirm optical clarity and demonstrate compatibility of the grooved hydrogels produced with the ligHTS method with a complementary imaging technique, which can increase the throughput of quality control of the hydrogel geometry beyond confocal fluorescent z-stacking.

## 3. Discussion

We developed a plate-wide, absorber-guided photopolymerization approach that overcomes a key bottleneck in imaging-based UHTS by enabling uniform, meniscus-free, anisotropic GelMA hydrogels across 384/1536-well plates using collimated light.

Tartrazine-mediated axial attenuation translates UV dose into predictable gel thickness and surface relief, allowing full-plate patterning without focusing optics. The resulting hydrogels preserve optical clarity and native-like stiffness, capturing canonical mechano-phenotypes such as reduced spreading on soft matrices and contact-guided migration (Fig. 2e-i, Fig. 3d-f, Fig. 4b-h, Fig. 5c-g, Fig. 6a-f).

In contrast to previous methods limited to rigid or 96-well plate formats that required molding or bottomless assemblies, LigHTS generates spacer-free microtopographies through a single collimated exposure using a film photomask. Continuous grooves are patterned across all wells up to1536-well plates with high pitch fidelity and ±20% coefficient of variation in thickness and depth, satisfying HTS geometric standards while eliminating optics, spacers, and manual alignment (Fig. 5c-g, Fig. 6a-c). To our knowledge, this represents the first demonstration of microgrooved hydrogel substrates in 384/1536-well formats,^34–37,41–43,47,53^ establishing LigHTS as a scalable route for high-density, mechanically and architecturally tunable biomaterial screening. As shown in Figure 5b, this fabrication method does not require ultra-high light collimation, and can be performed using a sub-10$ nail polish oven, eliminating any infrastructure costs. As a result, the cost per well is mainly dominated by the price of the required volume and concentration of the GelMA solution (around $0.4 and $0.1 per 384-wells and 1536-well plates, respectively), which is in the order of magnitude compatible with large screening price requirements. The realized hydrogel functionalization supports high-NA imaging for single-cell phenotypes and holographic phase imaging for rapid quality control, shortening feedback loops between fabrication and assay while retaining mechanobiological relevance for aligned tissues such as nerve, muscle, and fibrous stroma (Fig. 3d-f, Fig. 5e-g, Fig. 6a-d).^8,28–30^ This creates a practical path to deploy stiffness- and alignment-aware readouts in discovery campaigns at scale.

While the current UV-based process limits axial feature complexity and is not yet cell-compatible, transitioning to longer-wavelength initiators and optimized hydrogel precursors could mitigate phototoxicity, enabling direct on-cell patterning. Similarly, extending from the currently proposed 2.5D geometry and single cell line to 3D niches for organoid seeding, in-plate encapsulation and, after absorber removal, multi-material pattern could broaden biological scope to barrier-like structures and multi-cell models.^13–15,22,45^ Technique robustness is currently limited to in-plate geometry and behavior, while between-plate stability and capability indices still need quantification. The process could therefore be hardened by implementing plate-level irradiance mapping, closed-loop dose compensation, and holographic-to-confocal calibration to ensure between-plate uniformity and regulatory compliance. Establishing process consistency and quality control parameters would also enhance translational capabilities across formulations. In fact, since the absorber-controlled working curve follows Beer-Lambert kinetics, the LigHTS principle generalizes readily to diverse photopolymerizable materials, including ECM-derived scaffolds, synthetic materials, and emerging photoactive mixed ionic-electronic conductors (OMIECs), enabling broader applicability beyond GelMA while retaining the fabrication and quality control advantages demonstrated here (Fig. 2g-i, Fig. 4b-h).^45,47^ This could open a path to cleanroom-free, all-optical fabrication of scalable biointerfaces for tissue modeling, sensing, and drug discovery. Collectively, these attributes position LigHTS as a practical, extensible, and cost-effective foundation for mechanobiology-aware screening platforms.

## 4. Experimental Section

*Wellplate cleaning and silanization*: to promote GelMA adhesion, glass-bottom 384-well plates (P384-1.5H-N, Cellvis) or 1536-well plates (M4562, Sigma-Aldrich) were cleaned and silanized before casting. Wells were incubated 1 h at 37 °C with 2% v/v Hellmanex™ III (Z805939, Sigma-Aldrich) in sterile-filtered DI water, rinsed 3× with 37 °C DI water, and dried. The glass bottom of the wells was then silanized with 3-(trimethoxysilyl)propyl methacrylate (TMSPM, 44059, Sigma-Aldrich): 0.4% v/v TMSPM in aqueous solution at pH 3.5 (glacial acetic acid, 33209-M, Sigma-Aldrich), stirred until clear; wells were incubated 1 h at RT, rinsed 2× with 37 °C DI water and once with ethanol, then dried. Plates were used immediately or sealed (Parafilm, 11772644, Fisher Scientific) and stored RT ≤ 1 week.

*Preparation of the hydrogel precursor solutions:* for testing the effect of different tartrazine concentration on gel thickness, a stock of 100 mg mL⁻¹ GelMA with 2.5 mg mL⁻¹ LAP in PBS (D16110025360, Cellink) was mixed with tartrazine (T0388, Sigma-Aldrich), LAP (900889, Sigma-Aldrich), and glycerol (327255000, Thermo Fisher Scientific) in PBS (pH 7.4; 10010015, Gibco) to obtain 75 mg mL⁻¹ GelMA, 2.5 mg mL⁻¹ LAP, 10% v/v glycerol, and tartrazine 0-9 mg mL⁻¹.

For testing the effect of different concentration of GelMA on scaffold thickness and stiffness, groove generation, and cell response, lyophilized GelMA (Type A, Bloom 300; degree of methacrylation > 75%) from Advanced Biomatrix (5208) or Sigma-Aldrich (938408) was dissolved in PBS. LAP and tartrazine were pre-dissolved in PBS to target 1-5 mg mL⁻¹ (LAP) and 2.25-9 mg mL⁻¹ (tartrazine), stirred until clear, sterile-filtered (0.22-µm pore PVDF membranes), and used immediately or stored 4 °C, protected from light, ≤ 4 days. To dissolve GelMA efficiently, the LAP/tartrazine solution was heated to 45-60 °C. To increase viscosity and decrease oxygen solubility, glycerol was added to reach 10% v/v of the hydrogel precursor solution. Then GelMA was added the solution to reach a concentration in the range 50-200 mg mL⁻¹ and stirred at 1000 rpm ≥ 20 min until fully dissolved.

Specifically, for testing contact guidance, lyophilized GelMA was dissolved in PBS containing 1 mg mL⁻¹ LAP and 4.5 mg mL⁻¹ tartrazine to reach the concentration of 150 mg mL⁻¹.

*Casting of the hydrogel precursor solution in multiwell plates:* to avoid physical GelMA gelation during casting, the receiving well plates and pipette tips were heated up to keep the viscosity of GelMA low during casting. Silanized plates were equilibrated 20 min at 37 °C. For 384-well plates, 100 µL tips pre-warmed in 45-60 °C DI water transferred ≥ 20 µL precursor per square well (≈ 11 mm² area). For 1536-well plates, 12.5 µL tips transferred ≥ 5 µL (≈ 2.3 mm² area). Plates were centrifuged 5 min at 1500 G (SL16R, Thermo Fisher Scientific) to remove bubbles and wet the glass. Plates were cooled 5 min at 4-8 °C, then exposed immediately or within 2 h (protected from light).

*DMD photopatterning, development, and post-exposure UV flood for flat gels*: A PRIMO maskless DMD system (Alvéole) on a Nikon ECLIPSE Ti2, controlled by Leonardo v2.4 (Alvéole), illuminated through a 4× air objective (CFI Plan Fluor 4×, NA 0.13, MRH00045). The single-field area was ≈ 2.4 mm × 1.4 mm. Irradiance at 365 nm at full-field 100% was ≈ 0.9 mW mm⁻² (estimated from 3 mW measured with Argo-POWER HM Slide V2, Argolight). Uniform exposures used Leonardo dose control to reach 4-20 mJ mm⁻². Development used warm (37 °C) MES buffer (25 mM, pH 6.0 ± 0.1; 76039, Sigma-Aldrich), prepared from 0.5 M stock in DI water with NaOH (S2770) pH correction. Gels were incubated in MES three times with 30 min exchanges at 37 °C. After the third rinse, gels were left overnight at 37 °C, then rinsed once more. For the post-exposure flood, LAP (1 mg mL⁻¹) in MES was added (20 µL in 384-well; 5 µL in 1536-well) and plates were flood-exposed using a commercial UV LED nail lamp (365 nm, > 10 mW cm⁻²) for 15 min. Gels were rinsed 3× with PBS and stored in PBS containing 0.2% v/v Primocin (ant-pm-05, InvivoGen) ≥ 24 h before cell seeding.

*Gel surface functionalization using fluorescent beads*: To label the gel-liquid interface for profilometry, surfaces were coated at RT using MES throughout. Samples were equilibrated once in MES; poly-L-lysine (PLL, P4707, Sigma-Aldrich) 0.0025% w/v in MES (from 0.01% w/v stock) was applied 30 min, then rinsed once with MES (2 min). Carboxylated polystyrene beads (FluoSpheres™, 0.2 µm, 660/680 nm, F8807, Invitrogen; 2% solids) were freshly diluted 1:100 in MES, dispensed, and plates were centrifuged 5 min at 500 × g to accelerate sedimentation, followed by 10 min static incubation. After gentle aspiration and one MES rinse, a PLL “cap” (0.0025% w/v in MES) was applied 15 min and rinsed twice with MES. PBS was then added to half-volume to equilibrate osmolarity; 15 min incubation at 37 °C; finally, wells were fully exchanged to PBS and equilibrated ≥ 1 h at 37 °C before imaging.

*Gel profile characterization by confocal fluorescence microscopy:* Spinning-disk confocal (Crest V3 X-Light on Nikon) with CELESTA Light Engine (TSX5030FV, Lumencor) and a Photometrics Kinetix sCMOS camera acquired 4×4-binned z-stacks of the bead layer. Objectives and z-sampling were chosen by expected thickness: 4× air (CFI Plan Fluor 4×, NA 0.13) for thick gels (> 1 mm) and quick screening, Δz = 30 µm; 10× air (CFI Plan Apo 10×, NA 0.30, WD 1.6 mm, MRH00105), Δz = 5.6 µm, 3×3 tiles; 20× air (CFI Plan Apo 20× λ, NA 0.75, MRD00205), Δz = 0.9 µm, 6×6 tiles. Reference beads on the glass (same well or neighbor wells) set the zero plane and allowed tilt correction.

*Z-stack analysis of gel thickness*: Z-stacks were processed with a custom Python pipeline (see GitHub for details). Z-stacks were converted to per-pixel height maps by sub-pixel quadratic refinement at the axial intensity peak. A local lower-envelope was computed by grey-scale erosion with a disk of radius r_px chosen adaptively from the nominal bead diameter and pixel size. A least-squares plane was fit on the lower-envelope within the ROI and only the tilt terms were subtracted. An ISO 16610 Gaussian S-L band-pass was then applied with λs set adaptively as λs = max(2.5 × d_eff, 2.355 × σ_min_px × Δx) and λc = 120 µm. Inliers for statistics were defined on the L-filtered surface using a 1-99% percentile window. We report Sa, Sq, Sz, Ssk, Sku, residual Sq, and flatness by PV, p95-p05, and minimum-zone. For truncated stacks (≥30% of argmax at first/last slice), a strict truncated path reports absolute heights without salvage extrapolation. The ROI was exceeding the well size for all analyses for Figure 1h and was smaller (around 2 mm x 2 mm within the gel) for Figure 1i.

*Culture of HT1080 fibrosarcoma cell line:* Human fibrosarcoma cell lines (HT1080) were previously engineered to stably express the tubulin reporter RFP in addition to the F-actin reporter LifeAct-EGFP established in the commercial line (HT-1080-LifeAct-TagGFP2, 40101, Ibidi). A genetically-encoded variant of a FUCCI cell cycle sensor with CFP and iRFP signals for G1 and S/G2/M phases was also present in the cells, but not used in this study. The cells were cultured using complete cell culture media, consisting of DMEM / F-12 with L-glutamine, HEPES, and phenol Red (11330032, Gibco), 10% v/v heat-inactivated fetal bovine serum (FBS, 10270106, Gibco), and 1% v/v Penicillin-Streptomycin (10,000 U mL⁻¹, 15140-122, Gibco), and passaged using Trypsin-EDTA solution (T4049, Sigma-Aldrich) before the cells could reach 90% confluency. To prevent remaining trypsin during cell passaging from digesting the GelMA hydrogels, the cell suspensions were always centrifuged and resuspended in trypsin-free media.

*Gel compatibility with with high resolution imaging:* to verify that the photopatterned gels are optically clear and that the thickness is small enough to allow high-resolution imaging with standard work distances (< 400 μm), HT1080 cells were seeded on the gels and imaged after two days of culture using a Crest V3 X-Light spinning disk confocal microscope (Nikon) equipped with a Celesta Light Engine source (TSX5030FV, Lumencore), a Photometrics Kinetix Scientific CMOS camera, and a 100x silicon oil objective (CFI SR HP Plan Apo Lambda S 100XC Sil, N.A. 1.35, WD 0.31 mm, MRD73950, Nikon). Specifically, images were acquired of actin and tubulin by laser excitation at 477 nm and 546 nm respectively, no binning, 1s and 200 ms exposure time respectively.

*Mechanical characterization of the gel properties:* to verify the possibility to tune the stiffness of photocrosslinked GelMA and produce soft (<10 kPa) gels, large (>1 cm diameter) gels with GelMA concentration from 50 to 200 mg mL⁻¹ were produced on 2-cm diameter glass coverglass (631-1343, Avantor) adapting the procedure described for the gel patterned in wellplates and illuminate uniformly to reach 15 mJ mm^−2^. After development and UV flooding, the gels were incubated in 2% w/v BSA in PBS for 4 hours, and rinsed twice with PBS.

Nanoindentation mechanical characterization was performed using a Chiaro nanoindenter (Optics11 Life, Amsterdam, The Netherlands). Specifically, a probe comprising a reflective cantilever with stiffness of 0.48 N/m and a spherical tip with radius 25 µm was used to perform gel indentation of 5 µm to extract the elastic modulus and evaluate the viscoelastic properties of the materials. The effective Young’s modulus obtained from a Hertzian sphere-on-half-space was fed to a custom-made Python script (3.11; pandas 1.5.3, NumPy 1.24.0, SciPy 1.14.1, statsmodels 0.13.5, matplotlib 3.6.3, seaborn 0.11.2) to perform the statistical analysis. We treated indentations as technical replicates nested within experimental replicates. Planned pairwise comparisons were performed with two-sided Mann-Whitney U tests, followed by Holm step-down correction to control the family-wise error rate at α=0.05. For each contrast we report the U statistic, Holm-adjusted p value, and significance call.

*Cell area extraction:* to automate cell segmentation and cell area quantitative analysis a custom Python pipeline was developed. Denoised image stacks of different experiments were placed into separate folders, which were processed using the new Cellpose-SAM model to jointly segment tubulin and actin channels,^54^ yielding reliable whole-cell masks. For each acquisition, the script generates side-by-side plots of raw images and segmentation masks, saves the masks as TIFF files, and extracts single-cell features with *scikit-image*, storing cell areas in csv files within dedicated subdirectories. A second script parses csv outputs and generates histograms and overview plots of cell area distributions. Finally, a merging script concatenates all csv files across experiments, annotates files with experimental conditions, standardizes field-of-view indices, and produces a single dataset suitable for downstream statistical analysis.

*Statistical analysis general algorithm:* on large datasets (N>30) statistical analysis was performed through a custom Python script that, given the independent and dependent variables, checks the dataset using Levene’s test for homogeneity of variance and Shapiro-Wilk on model residuals. If assumptions hold, it runs a one-way ANOVA followed by Tukey’s HSD post-hoc test for comparison of the means between groups; otherwise, it applies Kruskal-Wallis with Dunn’s test (Holm-adjusted), or Mann-Whitney U tests. The workflow produces swarm plots with significance bars, and exports a docx report summarizing descriptives, assumption tests, performed tests and post-hoc results.

*Photopatterning, development, and UV flooding of the grooved GelMA hydrogels:* to produce grooved gels, the same methodology described for the preparation of the hydrogel precursor solution for flat GelMA was adopted. To generate a structured surface, a gradient illumination pattern was designed in Affinity Designer 2 consisting in a grayscale line array pattern matching the pixel count of PRIMO with intensities varying between the 30% and the 100% of the nominal UV dose, and then exposed at 10 mJ mm^−2^. The illumination was performed in grayscale mode with Leonardo software tuning the exposure of the individual areas to reach the intended local dose. To map geometry control over grooved gels, different gradient illumination patterns were designed in Affinity Designer 2 as described, varying the line widths and spacing (20-28-40 μm in this study) and then exposed in PRIMO through Leonardo software with different selected local doses between 5 and 21 mJ mm⁻². For contact guidance experiments, the grayscale lined pattern consisted of features of 20 μm width with 20 μm spacing. The gradient grayscale values corresponded to 30% to 100% of the selected local illumination of the tool, that was fixed at 8 mJ mm⁻².

*Gel surface coating and imaging:* to perform a qualitative check of proof-of-concept surface patterning results, hydrogel surface was incubated in 20 μg mL-1 FITC-tagged fibronectin (FNR01-A, Sigma-Aldrich) in PBS without Ca²⁺/Mg²⁺ (PBS -/-) for 30 minutes, then rinsed three times with PBS -/-to remove excess coating. We used a Crest V3 X-Light spinning disk confocal microscope (Nikon) equipped with a Celesta Light Engine source (TSX5030FV, Lumencore), a Photometrics Kinetix Scientific CMOS camera, and a 20 × air objective (CFI Plan Apo 20x Lambda, N.A. 0.75, MRD00205, Nikon) to image the gel-liquid interface. Specifically, we acquired a stack using confocal mode, 477 nm light source, 4×4 binning, 10 ms exposure time, and volume was reconstructed from the stack with the integrated NIS Element software.

*Image analysis for geometric characterization:* To quantify surface patterning, we coated the gel surface with fluorescent beads and imaged z-stacks as described above (Gel profile characterization by confocal fluorescence microscopy). A Python routine converted each stack into a height map. For every pixel it found the z-slice of maximum intensity, refined the position with a three-point quadratic vertex (ISO 5436-2 Type F), and multiplied by Δz to obtain micrometre-level heights. Noise was reduced with a mirrored 5-px median and a k = 5×MAD spike filter, a least-squares plane was removed (ISO 25178-3), then a mild 3×3 median was applied. A 1st-99th percentile mask excluded bright aggregates. For groove alignment, the 2-D FFT energy was scanned over angles. If the dominant orientation deviated by more than 1°, the map was rotated to that angle and cropped to the maximal inscribed rectangle to avoid edge effects. Pitch was derived in two ways. Spectrally, the average-profile FFT gave the dominant frequency fₘ and a first estimate P = 1/fₘ. Spatially, each row was low-pass filtered with a zero-phase Gaussian cut-off at 0.8×P, peaks were detected using spacing and prominence rules seeded by the FFT estimate, and interquartile outliers were removed. The robust mean and standard deviation of the remaining spacings were reported as pitch and spread. Depth was computed using lightly smoothed profiles to lock crest and valley positions while reading amplitudes on unfiltered data to avoid bias. Quadratic sub-slice extrema refined positions. No scalar amplitude correction was applied. Uncertainty from lateral sampling (pixel pitch/√12), axial sampling (Δz/√12), and a relative calibration term of 1.2% of depth was combined in quadrature and reported. Repeated-scan data were analysed per AIAG MSA repeatability and reproducibility. Results exceeding 10% of tolerance were flagged and the percentage logged for ISO/TS 22514 use. Capability indices Cp and Cpk were computed against predefined ±10% design specifications, with a pass criterion of ≥1.33 (ISO/TS 22514-4). Depth distributions were also monitored with three-sigma control limits. A separable 2-D Gaussian (λₛ equal to the pixel pitch) separated roughness from waviness. Areal Sa and Sq (ISO 25178-2) and profile Ra, Rq, and Rz (ISO 4287) were reported. All per-sample results were saved to CSV and used for the statistical analysis described in Statistical analysis—general algorithm.

*Contact guidance long term live imaging experiments:* to verify that the hydrogel surface anisotropies were able to provide contact guidance to cells, HT1080 cells were seeded on LigHTS-produced grooved gels and flat gels, and on glass pre-coated with 40 µg mL⁻¹ human plasma fibronectin (F0895, Sigma-Aldrich). Cell migration was observed with a 12-hour live imaging experiment. Specifically, the cells were cultured as described before, then 48 hours before the migration assay, HT1080-containing trypsin-free media was added to each well to seed roughly 1000 cells per well in the 384-well plate format and roughly 250 cells per well in the 1536-well plate format. Cells were let grow in the incubator until the start of the experiment. To acquire the 12-hour migration experiments, a Crest V3 X-Light spinning disk confocal microscope (Nikon) equipped with a Celesta Light Engine source (TSX5030FV, Lumencore), a Photometrics Kinetix Scientific CMOS camera, and a 20 × air objective (CFI Plan Apo 20x Lambda, N.A. 0.75, MRD00205, Nikon) was used. In terms of imaging conditions, the tubulin and actin signal in confocal mode and the transmitted white spectrum light in differential interference contrast (DIC) mode were recorded in a 15-minute-interval time series for 12 hours. Specifically, tubulin was imaged with excitation at 546 nm, 4×4 binning, 200 ms exposure, actin was imaged with excitation at 477 nm, 4×4 binning, 50 ms exposure, and DIC was imaged in 4×4 binning, 4 ms exposure. The fields of view that showed consistent focus through the 12 hours and no mechanical failure in gels were successively analyzed.

*Substrate-dependent analysis of HT1080 migrations:* to analyze migratory behaviour of HT1080 cell line on different substrate, the confocal-acquired raw time-stacks were denoised using NIS-Elements “Denoise” feature and analysed as follows. First, the images were converted from the Nikon proprietary .nd2 format to .tiff files and the channels were split for further analysis through an ImageJ macro. The .tiff file containing only the tubulin channel was further processed through custom Python scripts. The first analysis script uses CellPose 3.1.0 pretrained cyto3 network to automatically segment the stacks given the user-selected mean cell diameter,^55^ corresponding to 17 pixels in these experiments. The labels are imported in LapTrack (v 0.17.0) to generate tracks describing cell motion,^56^ tracks are saved in a .csv file for later use. To be able to visually check the cell segmentation and track attribution, a .mov movie overlaying the cell label border and its assigned track to the tubulin signal is created through matplotlib and saved.

The tracks contained in the .csv files are then used for quantitative analysis of cell movement on different substrates and between three independent experiments via another custom Python script. For each experiment, the script reads the tracks and calculates the parameter used to determine the contact guidance of grooves, called “2D nematic order parameter”. This parameter measures the deviation of the angle between the starting and ending point of the track segment (θ) from the vertical direction that corresponds to grooves orientation: *NO* = *cos* (2 ∗ (*θ* − 90°)). By construction, this parameter varies between -1 if the cells migrate exclusively in the direction perpendicular to the grooves to 1 if all the movements are perfectly parallel to the grooves. Then, the script assigns a weighted nematic order (WNO) value to each track in the field of view, weighting the value of the single displacement with respect to its length. A synthetic WNO of the entire field of view is calculated, weighting again the WNO per track on its length. This parameter is used to compare each field of view with others recorded during the same experiment and among different experiments via a permutation test. In addition to the WNO, the code also calculates and compares the mean cell velocity of the field of view. All analysis results are reported in a .csv file, used to perform statistical analysis.

To perform statistical analysis, a custom Python routine is used. The script asks the user to select a .csv file containing a row per field-of-view (FOV) measurements across experimental groups. After user-defined selection of dependent variable, group labels, experiment identifiers, and optional FOV identifiers via a graphical interface, values are aggregated to yield one entry per experiment × group × FOV. Due to limited replication of experiments (N=3, with N=3 technical replicates per group per experiment) statistical testing relies on exact blocked label-permutation, in which group labels were exhaustively permuted within each experiment while preserving sample sizes; the test statistic was the mean difference of group means across experiments, and two-sided p-values with 95% permutation confidence intervals were obtained. P-values were Holm-adjusted where applicable. The code generates a dependent variable vs groups plot highlighting the statistical difference among groups and a docx report of the statistical analysis performed.

*Optimization of hydrogel preparation and photopatterning for parallel processing:* to optimize the hydrogel composition and photopatterning procedure for upright large-scale illumination produced by large-scale highly collimated light sources present in micromachining mask aligners, a test setup for multiwell photopatterning was assembled by using the previously mentioned commercial UV nail polish oven. Specifically, all but the square LED arrays in the center of the module projecting 355 nm light vertically on the sample were blocking. The power output was measured with Argo-POWERHM Slide V2 and was stable at around 5 mW cm^−2^ in a 5 x 5 cm^2^ below the square LED array. Due to the upright illumination, the plate had to be inverted during exposure. To ensure that the hydrogel precursor solution stayed in place, 10% v/v glycerol (Glycerol, 99.5%, 327255000, Thermo Scientific Chemicals) was added to the solution to increase the viscosity. Moreover, the LAP content was increased to 5 mg mL⁻¹ to avoid polymerization-induced phase separation (PIPS) associated with the significantly lower irradiance in this configuration compared to Primo DMD (20-fold reduction). No modifications were made to the previously described procedures for gel development and UV flooding.

For parallel processing, the light source of a Karl Suss MA6 with I-line filter was used (365 nm cutoff, mercury lamp 3.5 mW cm^−2^ power at 365 nm) A whole plate film photomask (JD photodata) was manually aligned to the wells and put in contact with a quartz block. The UV exposure was carried out in contact with the quartz for 16 minutes. No modifications were made to the previously described procedures for gel development and UV flooding. For parallel produced gel characterization, the same procedures of fluorescent beads coating, confocal imaging, and quantitative analysis were carried out as described above.

*Large-scale compatible characterization of groove profile by holographic phase imaging:* to efficiently characterize the gel profile in a large number of wells, holographic phase imaging using HoloMonitor M4 (Phase Holographic Imaging AB) was used to characterize the gel profile in the 1536-well plate format. The measurement was performed using a 20x air objective (Olympus PLN 20X, NA 0.4, N1215900, Evident), a low-power laser unit (635 nm, 0.2 mW cm^−2^). The plate top lid was removed and the wells were filled (around 100 μL) with PBS containing Ca²⁺/Mg²⁺ (PBS +/+) to provide physiological osmolarity and minimize the curvature of the air-water interface which would otherwise disturb the measurement. The resulting grayscale PNG images were analysed using a dedicated Python script. Specifically, the code converts the grayscale height-map of the grooved surface into micron units, removes slow background curvature with a large-sigma (50) Gaussian filter, and traces the mean height of every image row to detect ridge and valley peaks. It then selects only ridge-valley pairs that are at least 5 pixels apart and ≥ 4 µm deep, then quantifies groove uniformity with two coefficients of variation (CV). “Depth CV” is the relative standard deviation of the ridge-to-valley depths, while “Shape CV” is derived by resampling each groove profile to 100 points, detrending and z-scaling it, computing all pairwise Euclidean distances between normalized profiles, and taking the CV of those mean distances. These metrics capture, respectively, how much the grooves vary in depth and in cross-sectional shape across the image. The points spread is compared between groups with different nominal pitch and against confocal characterization for validation purposes.

## Acknowledgements

A.E. and S.R. contributed equally to this work. The authors would like to acknowledge that this project has received funding from the European Union’s Horizon 2020 research and innovation programme under the Marie Sklodowska-Curie grant (HORIZON-MSCA-2023-PF, agreement No 101153603) awarded to A.E. This work was also supported by the European Research Council (ERC) Starting Grant No. 852560 to F.S.P, as well as by the Italian Ministry of Education, University, and Research (MIUR) (FARE2020 grant, No. R20ZE54CTK) to F.S.P. This project is supported by the Chips Joint Undertaking (Grant Agreement No. 101140192 to FSP) and its members, including the top-up funding of Belgium, Germany, Hungary, Ireland, Italy, the Netherlands, Portugal, Romania, and Spain. For Italy, the work was funded by the Italian Ministry of Enterprises and Made in Italy (MIMIT) under CUP F13C23003260003. While the work was supported by the European Union, the views and opinions expressed are those of the author(s) only and do not necessarily reflect those of the European Union or Chips Joint Undertaking. Neither the European Union nor the granting authority can be held responsible for them.

The authors also thank Prof. Frank Niklaus for providing the film photomask and Fabio De Ferrari and Cecilia Aronsson for their assistance in the proof of concept experiment with collimated light gel exposure. This work was partially performed at PoliFAB, the microtechnology and nanotechnology center of the Politecnico di Milano, and the authors thank Chiara Nava for the support with the collimated light exposure setup. The authors also thank Prof. Michele Conti for the access to the nanoindentometer and Giulia Formenton for the tool training.

## References

1. Paul, S. M. et al. How to improve R&D productivity: the pharmaceutical industry’s grand challenge. Nat Rev Drug Discov 9, 203–214 (2010).

2. Schuhmacher, A., Hinder, M., von Stegmann und Stein, A., Hartl, D. & Gassmann, O. Analysis of pharma R&D productivity – a new perspective needed. Drug Discovery Today 28, 103726 (2023).

3. Ringel, M. S. What is the right amount to spend on biopharma R&D? Nature Reviews Drug Discovery 16, 597–598 (2017).

4. Scannell, J. W., Blanckley, A., Boldon, H. & Warrington, B. Diagnosing the decline in pharmaceutical R&D efficiency. Nat Rev Drug Discov 11, 191–200 (2012).

5. Sun, D., Gao, W., Hu, H. & Zhou, S. Why 90% of clinical drug development fails and how to improve it? Acta Pharmaceutica Sinica B 12, 3049–3062 (2022).

6. Xie, R. et al. A comprehensive review on 3D tissue models: Biofabrication technologies and preclinical applications. Biomaterials 304, 122408 (2024).

7. Foglietta, F., Canaparo, R., Muccioli, G., Terreno, E. & Serpe, L. Methodological aspects and pharmacological applications of three-dimensional cancer cell cultures and organoids. Life Sciences 254, 117784 (2020).

8. Honarnejad, S., van Boeckel, S., van den Hurk, H. & van Helden, S. Hit Discovery for Public Target Programs in the European Lead Factory: Experiences and Output from Assay Development and Ultra-High-Throughput Screening. SLAS Discovery 26, 192–204 (2021).

9. Kwon, K. K., Lee, J., Kim, H., Lee, D.-H. & Lee, S.-G. Advancing high-throughput screening systems for synthetic biology and biofoundry. Current Opinion in Systems Biology 37, 100487 (2024).

10. Chen, W. L. K. & Simmons, C. A. Lessons from (patho)physiological tissue stiffness and their implications for drug screening, drug delivery and regenerative medicine. Advanced Drug Delivery Reviews 63, 269–276 (2011).

11. Peterson, N. C., Mahalingaiah, P. K., Fullerton, A. & Piazza, M. D. Application of microphysiological systems in biopharmaceutical research and development. Lab Chip 20, 697–708 (2020).

12. Inglese, J. et al. Quantitative high-throughput screening: A titration-based approach that efficiently identifies biological activities in large chemical libraries. Proceedings of the National Academy of Sciences 103, 11473–11478 (2006).

13. Rossi, G., Manfrin, A. & Lutolf, M. P. Progress and potential in organoid research. Nat Rev Genet 19, 671–687 (2018).

14. Low, L. A., Mummery, C., Berridge, B. R., Austin, C. P. & Tagle, D. A. Organs-on-chips: into the next decade. Nat Rev Drug Discov 20, 345–361 (2021).

15. Wang, L., Hu, D., Xu, J., Hu, J. & Wang, Y. Complex in vitro model: A transformative model in drug development and precision medicine. Clinical and Translational Science 17, e13695 (2024).

16. Eric, E.-O. HIGH-THROUGHTPUT SCREENING QUALITY CONTROL GENERAL GUIDELINES.

17. Madani, A., Alvarez, N., Park, S., Murugan, M. & Perlin, D. S. Rapid luminescence-based screening method for SARS-CoV-2 inhibitors discovery. SLAS Discovery 31, (2025).

18. Booij, T. H., Price, L. S. & Danen, E. H. J. 3D Cell-Based Assays for Drug Screens: Challenges in Imaging, Image Analysis, and High-Content Analysis. SLAS DISCOVERY: Advancing the Science of Drug Discovery 24, 615–627 (2019).

19. Dave, R., Pandey, K., Patel, R., Gour, N. & Bhatia, D. Leveraging 3D cell culture and AI technologies for next-generation drug discovery. Cell Biomaterials 1, 100050 (2025).

20. Maoz, B. M. et al. A linked organ-on-chip model of the human neurovascular unit reveals the metabolic coupling of endothelial and neuronal cells. Nat Biotechnol 36, 865–874 (2018).

21. Novak, R. et al. Robotic fluidic coupling and interrogation of multiple vascularized organ chips. Nat Biomed Eng 4, 407–420 (2020).

22. Ingber, D. E. Human organs-on-chips for disease modelling, drug development and personalized medicine. Nat Rev Genet 23, 467–491 (2022).

23. Organ-Chip Models. Emulate https://emulatebio.com/organ-chips/.

24. Nasiri, R. et al. Engineering biomimetic tissue barrier models on chips: From design and fabrication to applications in disease modeling and drug screening. Biomaterials 327, 123739 (2026).

25. Mirbagheri, M. et al. Advanced cell culture platforms: a growing quest for emulating natural tissues. Mater. Horiz. 6, 45–71 (2019).

26. Sun, Y., Deng, R., Ren, X., Zhang, K. & Li, J. 2D Gelatin Methacrylate Hydrogels with Tunable Stiffness for Investigating Cell Behaviors. ACS Appl. Bio Mater. 2, 570–576 (2019).

27. Buxboim, A., Rajagopal, K., Brown, A. E. X. & Discher, D. E. How deeply cells feel: methods for thin gels. J. Phys.: Condens. Matter 22, 194116 (2010).

28. Luciano, M. & Gabriele, S. Designing hydrogel dimensionality to investigate mechanobiology. Soft Matter 21, 4551–4572 (2025).

29. Wang, W. Y. et al. Extracellular matrix alignment dictates the organization of focal adhesions and directs uniaxial cell migration. APL Bioeng. 2, 046107 (2018).

30. Lu, K. et al. Biofabrication of aligned structures that guide cell orientation and applications in tissue engineering. Bio-des. Manuf. 4, 258–277 (2021).

31. Torchia, E. et al. HYDRA: Fabrication of cell culture HYDrogels by Robotic liquid handling Automation for high-throughput drug testing. 2024.12.22.629679 Preprint at 10.1101/2024.12.22.629679 (2025).

32. Advanced BioMatrix -CytoSoft^TM^ Imaging, 96-Well Plates (0.2 -64 kPa). https://advancedbiomatrix.com/cytosoft-96well.html.

33. Softwell 384 Glass. Matrigen https://store.matrigen.com/products/softwell-384-glass.

34. Lücker, P. B. et al. A microgroove patterned multiwell cell culture plate for high-throughput studies of cell alignment. Biotechnology and Bioengineering 111, 2537–2548 (2014).

35. Hu, J. et al. High-Throughput Mechanobiology Screening Platform Using Micro-and Nanotopography. Nano Lett. 16, 2198–2204 (2016).

36. Conner, A. A. et al. High-throughput analysis of topographical cues for the expansion of murine pluripotent stem cells. Nanotechnology 35, 455101 (2024).

37. van der Boon, T. A. B. et al. Well Plate Integrated Topography Gradient Screening Technology for Studying Cell-Surface Topography Interactions. Advanced Biosystems 4, 1900218 (2020).

38. Lei, R. et al. Multiwell Combinatorial Hydrogel Array for High-Throughput Analysis of Cell–ECM Interactions. ACS Biomater. Sci. Eng. 7, 2453–2465 (2021).

39. Rossy, T. et al. Leveraging microtopography to pattern multi-oriented muscle actuators. Biomater. Sci. 13, 2891–2907 (2025).

40. Mih, J. D. et al. A Multiwell Platform for Studying Stiffness-Dependent Cell Biology. PLOS ONE 6, e19929 (2011).

41. Harkness, T. et al. High-content imaging with micropatterned multiwell plates reveals influence of cell geometry and cytoskeleton on chromatin dynamics. Biotechnology Journal 10, 1555–1567 (2015).

42. Almeida, F. V. et al. High-Content Analysis of Cell Migration Dynamics within a Micropatterned Screening Platform. Advanced Biosystems 3, 1900011 (2019).

43. Chitsaz, D. & Kennedy, T. E. High-throughput microcontact printing of proteins in microwell cell culture plates. MethodsX 12, 102665 (2024).

44. Zhang, B. et al. GelMA micropattern enhances cardiomyocyte organization, maturation, and contraction via contact guidance. APL Bioeng. 8, 026108 (2024).

45. Skelton, M. L. et al. Modular Multiwell Viscoelastic Hydrogel Platform for Two- and Three-Dimensional Cell Culture Applications. ACS Biomater. Sci. Eng. 10, 3280–3292 (2024).

46. Bril, M. et al. Digital Photoinduced Topographical Microsculpting of Hydrogels. Advanced Materials Technologies 9, 2400721 (2024).

47. Li, Y. et al. Theoretical prediction and experimental validation of the digital light processing (DLP) working curve for photocurable materials. Additive Manufacturing 37, 101716 (2021).

48. Li, Y. et al. High-fidelity and high-efficiency additive manufacturing using tunable pre-curing digital light processing. Additive Manufacturing 30, 100889 (2019).

49. Pariskar, A., Sharma, P. K., Murty, U. S. & Banerjee, S. Effect of Tartrazine as Photoabsorber for Improved Printing Resolution of 3D Printed “Ghost Tablets”: Non-Erodible Inert Matrices. Journal of Pharmaceutical Sciences 112, 1020–1031 (2023).

50. Levato, R. et al. High-resolution lithographic biofabrication of hydrogels with complex microchannels from low-temperature-soluble gelatin bioresins. Materials Today Bio 12, 100162 (2021).

51. Pezzotti, M. et al. Vertically Integrated System for Tracking and Assessing cell-cycle aware phenotypes under confinement. 2025.09.25.678497 Preprint at 10.1101/2025.09.25.678497 (2025).

52. Sante, M. D. et al. CALIPERS: Cell cycle-aware live imaging for phenotyping experiments and regeneration studies. 2024.12.19.629259 Preprint at 10.1101/2024.12.19.629259 (2025).

53. SmartGel. 4Dcell https://www.4dcell.com/smartgel.

54. Pachitariu, M., Rariden, M. & Stringer, C. Cellpose-SAM: superhuman generalization for cellular segmentation. 2025.04.28.651001 Preprint at 10.1101/2025.04.28.651001 (2025).

55. Stringer, C. & Pachitariu, M. Cellpose3: one-click image restoration for improved cellular segmentation. Nat Methods 22, 592–599 (2025).

56. Fukai, Y. T. & Kawaguchi, K. LapTrack: linear assignment particle tracking with tunable metrics. Bioinformatics 39, btac799 (2023).

